# Biased evolutionary inferences from bulk tumor samples

**DOI:** 10.1101/089680

**Authors:** J.M. Alves, T. Prieto, D. Posada

## Abstract

It is generally agreed that tumors are composed of multiple cell clones defined by different somatic mutations. Characterizing the evolutionary mechanisms driving this intratumor genetic heterogeneity (ITH) is crucial to improve both cancer diagnosis and therapeutic strategies. For that purpose, recent ITH studies have focused on qualitative comparisons of mutational profiles derived from bulk sequencing of multiple tumor samples extracted from the same patient. Here, we show some examples where the naive use of bulk data in multiregional studies may lead to erroneous inferences of the evolutionary trajectories that underlie tumor progression, including biased timing of somatic mutations, spurious parallel mutation events, and/or incorrect chronological ordering of metastatic events. In addition, we analyze three real datasets to highlight how the use of bulk mutational profiles instead of inferred clones can lead to different conclusions about mutational recurrence and population structure.

## INTRODUCTION

Over the past decade, global sequencing efforts of cancer genomes have revealed that genetic intratumor heterogeneity (ITH) represents a common feature of many cancer types^1^, arising from the progressive accumulation of somatic mutations within malignant cells during cancer evolution^2^. As a result, single tumors typically comprise multiple genetically distinct subpopulations of cells (i.e., clones), which may vary with regards to growth, metastatic potential, and therapeutic resistance^2,3^. With the increasingly obvious clinical implications of ITH^4^, a great deal of attention is currently being directed towards exploring the complex patterns of clonal heterogeneity under an evolutionary framework, in order to resolve the genetic history underlying cancer progression^5-7^ and gain a wider understanding of the evolutionary mechanisms driving tumor diversification^8–11^. Indeed, following the pioneer work by Gerlinger *et al.*^12^, reporting a high degree of variation in the genetic composition of primary tumors and metastases as a consequence of divergent clonal evolution, a number of studies have focused on the spatial and temporal dynamics of tumorigenesis, taking advantage of next-generation sequencing (NGS) data obtained from bulk tissue samples extracted from multiple tumor regions within a single patient^10,13-16^.

However, although a variety of statistical algorithms exist to infer the clonal composition of tumors from bulk NGS data (see Beerenwinkel *et al.*^17^ for an exhaustive review), most multiregional sequencing studies still analyze the spatial patterns of clonal diversity by directly comparing mutational profiles (absence/presence of mutations) across samples. For example, making use of whole-exome bulk sequencing data initially derived from 23 evenly distributed samples in a cross-section of a hepatocellular carcinoma, Ling *et al.*^18^ identified a set of 35 somatic single-nucleotide variants (SNVs) that were subsequently used to genotype approximately 300 samples of the same tumor. Ancestral relationships among the sampled regions were inferred from their mutational profiles, and used to delineate clonal boundaries and reconstruct clonal genealogies. Consistent with previous findings^10^, the authors observed extensive genetic diversity between all regional samples, pointing to a limited role of selection, with patterns of genetic diversity suggesting the appearance of new subclones on the peripheries of tumors which tend to radiate outwards. Similarly, Zhao*et al.*^19^ applied a series of phylogenetic methods to whole-exome sequencing data derived from 40 cancer patients to address the origin of metastases. By analyzing the mutational profiles of primary tumors and associated metastases, the authors concluded that metastatic lineages often evolved non-linearly from the primary tissue, suggesting that metastases may originate stochastically from distinct clonal lineages within the primary tumor of a given patient. In order to trace the timing of such metastatic events, the authors further transformed the inferred regional trees into patient-specific chronograms, calibrated a molecular clock based on different clinical parameters, and suggested that most metastatic lineages appeared to differentiate at very early stages of cancer progression, usually prior to clinical detection.

Importantly, at the heart of these studies is the implicit assumption that tumor clones present in a tissue sample can be meaningfully summarised as the collection of mutations observed in that sample (i.e., the mutational profile) – or that only a single or dominant clone exists per sample that carries all mutations –, and that reliable evolutionary relationships can be inferred from such information. However, given the high levels of ITH expected in most tumors, this assumption is not justified and can lead to biased inferences regarding the evolutionary history of a tumor.

## IMPLICATIONS OF MUTATIONAL PROFILES FOR EVOLUTIONARY INFERENCE

### Mutational histories

Since most tumors consist of a genetically heterogeneous population of cells, mutational profiles of bulk tumor samples essentially reflect the set of somatic mutations present in a detectable fraction of the cells sampled, but not immediately the collection of clones present. The fundamental reason for this is that, in the absence of single-cell information, the precise combination of mutations that occur in any given clone is unknown, and a set of *n* mutations can represent 1, 2 or even *n* clones. In the presence of ITH, multiple clones are expected per bulk sample, hence the observed mutational profile might easily correspond to a “composite clone” that never existed. Consequently, the use of mutational profiles as units for evolutionary analysis can have important implications.

To illustrate this idea, consider the cancer patient shown in Fig. 1A, whose primary tumor harbors three genetically distinct cell clones A-C resulting from the accumulation of five somatic mutations. The three clones share some mutations (“true clonal sequences”) that reflect their common history from a common ancestor (“true clonal phylogenetic tree”). For simplicity, we assume that the tumor does not contain healthy cells (i.e., no contamination). Fig. 1B depicts a hypothetical multiregional sequencing study, where three spatially separated regions from the primary tumor have been sampled and sequenced. While the sampling scheme shows that all three clones have been captured, the proportion of each cell clone per sample varies, with sample I consisting entirely of cells belonging to clone A, sample II being composed of cells from clones B (80%) and C (20%), and sample III carrying cells from clones A (30%) and C (70%). Accordingly, only the mutational profile for sample I corresponds to a true clonal sequence (clone A), while the profiles obtained for the other two samples represent a composite of clones BC and AC, respectively. If we now build a maximum parsimony (MP) tree using these composite clones (see Gerlinger*et al.*^12^; Gerlinger*et al.*^14^; Hao*et al.*^16^; Ling*et al.*^18^), we would wrongly infer that mutations 1, 2 and 3 occurred in the most recent common ancestor (MRCA) of these samples, and that mutation 5 occurred before mutation 4. Importantly, these problems can be avoided if one realizes that the history of the samples is often not the same as the history of the clones. Indeed, in recent years multiple algorithms have been developed for the identification of clones from bulk tumor samples^17^, usually by clustering mutations with similar variant allele frequency (VAF) into single clones. In this example, the clustering algorithm implemented in Clomial^20^ perfectly identifies the true clones. A MP tree of these clonal sequences accurately recovers the true clonal history and the right order of mutations (Fig. 1C, **Supplementary Note**).

**Figure 1.**
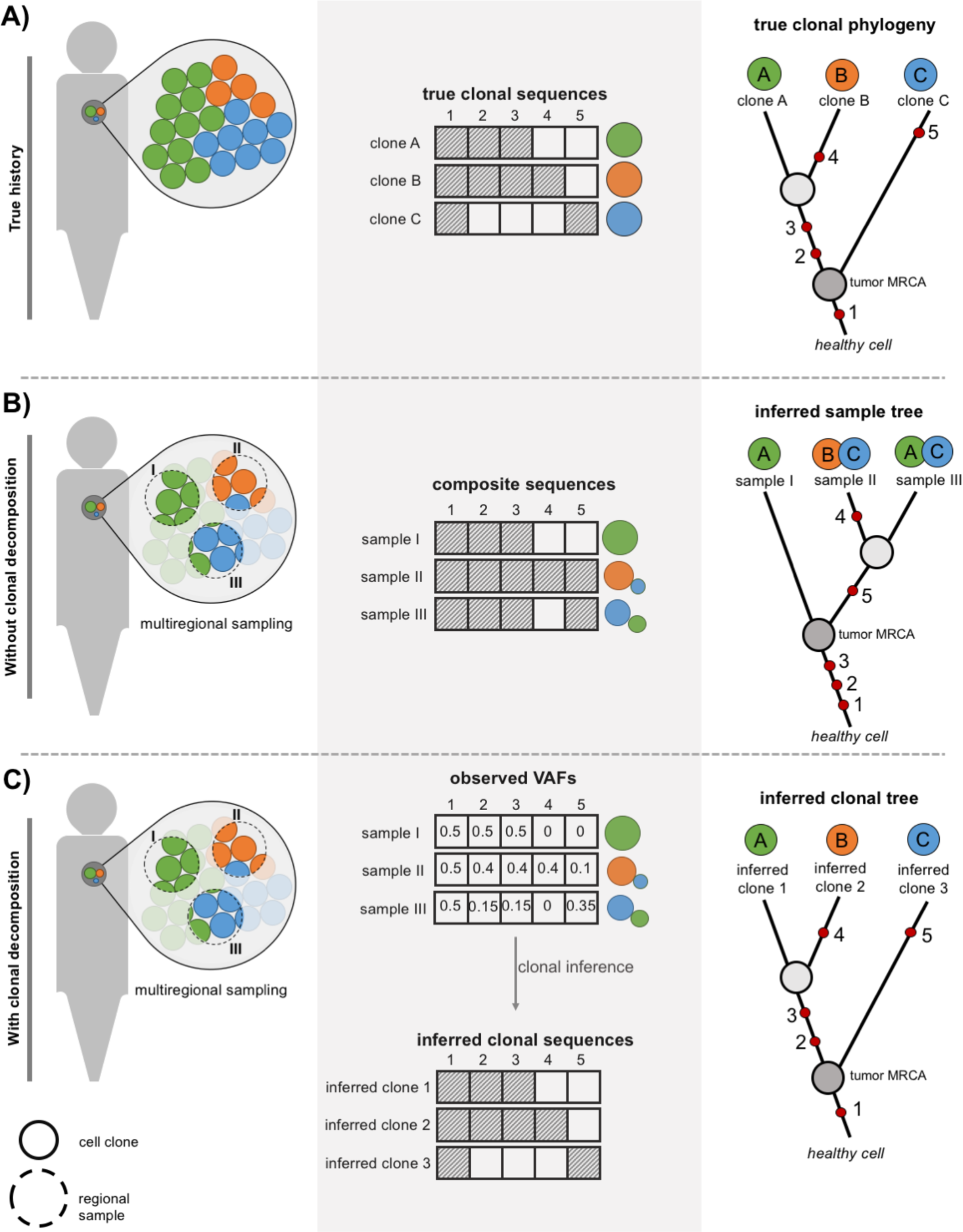
Phylogenetic analysis of bulk tumor samples (I). **A)** *Left-panel*: clonal composition of a hypothetical primary tumor. Colored circles represent the three clones present (A-C). *Mid-panel*: true clonal sequences for five different genomic sites, where the dashed square indicates a somatic mutation. *Right-panel*: true clonal history with red dots depicting the chronological order of mutations. Tumor most recent common ancestor (MRCA) highlighted as an internal node. **B)** *Left-panel*: bulk regional samples (I-III), with intermixed clones at different proportions. *Mid-panel*: composite sequences (presence/absence) inferred, dashed square indicates presence of mutation. *Right-panel*: inferred sample history using maximum parsimony. Red dots depict the inferred chronological order of mutations. **C)** *Left-panel*: bulk regional samples (I-III), with intermixed clones at different proportions. *Mid-panel*: variant allele frequency estimates for mutation at each sample, and inferred clonal sequences using the Clomial algorithm (see supplementary note for details*). Right-panel*: inferred clonal history using maximum parsimony. Red dots depict the inferred chronological order of mutations.

Fig. 2 illustrates another example involving the same hypothetical cancer patient (Fig. 2A), but for which a distinct set of regional samples was obtained (Fig. 2B). In this case, the use of mutational profiles results in a phylogenetic tree in which mutation 5 spuriously appears to have occurred twice, independently in sample II and III. Again, the use of a clustering algorithm for clonal identification avoids this type of bias, leading to the inference of the true tree and the true mutational history (Fig. 2C).

**Figure 2.**
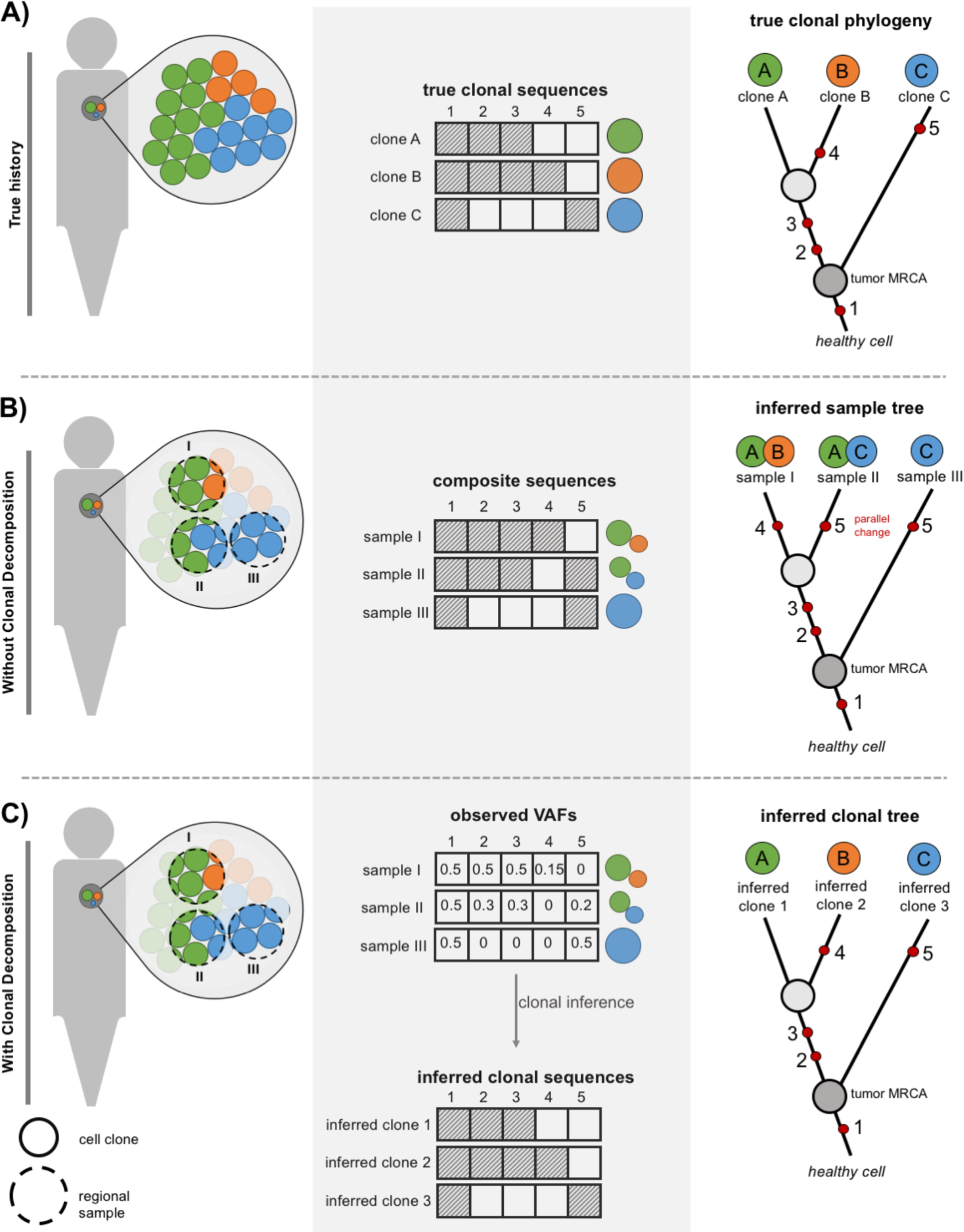
Phylogenetic analysis of bulk tumor samples (II). **A)** *Left-panel*: clonal composition of a hypothetical primary tumor. Colored circles represent the three clones present (A-C). *Mid-panel*: true clonal sequences for five different genomic sites, where the dashed square indicates a somatic mutation. *Right-panel*: true clonal history with red dots depicting the chronological order of mutations. Tumor most recent common ancestor (MRCA) highlighted as an internal node. **B)** *Left-panel*: bulk regional samples (I-III), with intermixed clones at different proportions. *Mid-panel*: composite sequences (presence/absence) inferred, dashed square indicates presence of mutation. *Right-panel*: inferred sample history using maximum parsimony. Red dots depicting the chronological order of mutations. **C)** *Left-panel*: bulk regional samples (I-III), with intermixed clones at different proportions. *Mid-panel*: variant allele frequency estimates for mutation at each sample, and inferred clonal sequences using the Clomial algorithm (but see supplementary note). *Right-panel*: inferred clonal history using maximum parsimony. Red dots depicting the chronological order of mutations.

Furthermore, the fact that different sets of samples obtained from the same primary tumor can generate two distinct, and incorrect, evolutionary histories (Fig. 1B and Fig. 2B) suggests that phylogenetic analysis of mutational profiles from bulk tumor tissues can be less straightforward than previously thought.

### Relative timing of metastasis

Another potential issue associated with the use of composite clones is determining the evolutionary relationships between the primary tumor and distant metastases. Following a similar approach to Zhao*et al.*^19^, consider now a patient for which four distinct samples have been sequenced: a primary tumor sample and three metastases (Fig. 3A). For simplicity, we assume that (i) there is no contamination from healthy cells, (ii) only the primary tumor hosts several clones, and (iii) somatic mutations accumulate linearly with time (i.e., following a molecular clock). In this example, there are four true clones (A-D). Clone A represents the ancestral lineage from which the other clones derived. Clone B, which was never sampled/existed in the primary tumor, represents the first metastasis (metastasis I), followed by migration of clone C (metastasis II) and later of clone D (metastasis III) into three distinct anatomical regions (Fig. 3B, right-panel).

**Figure 3.**
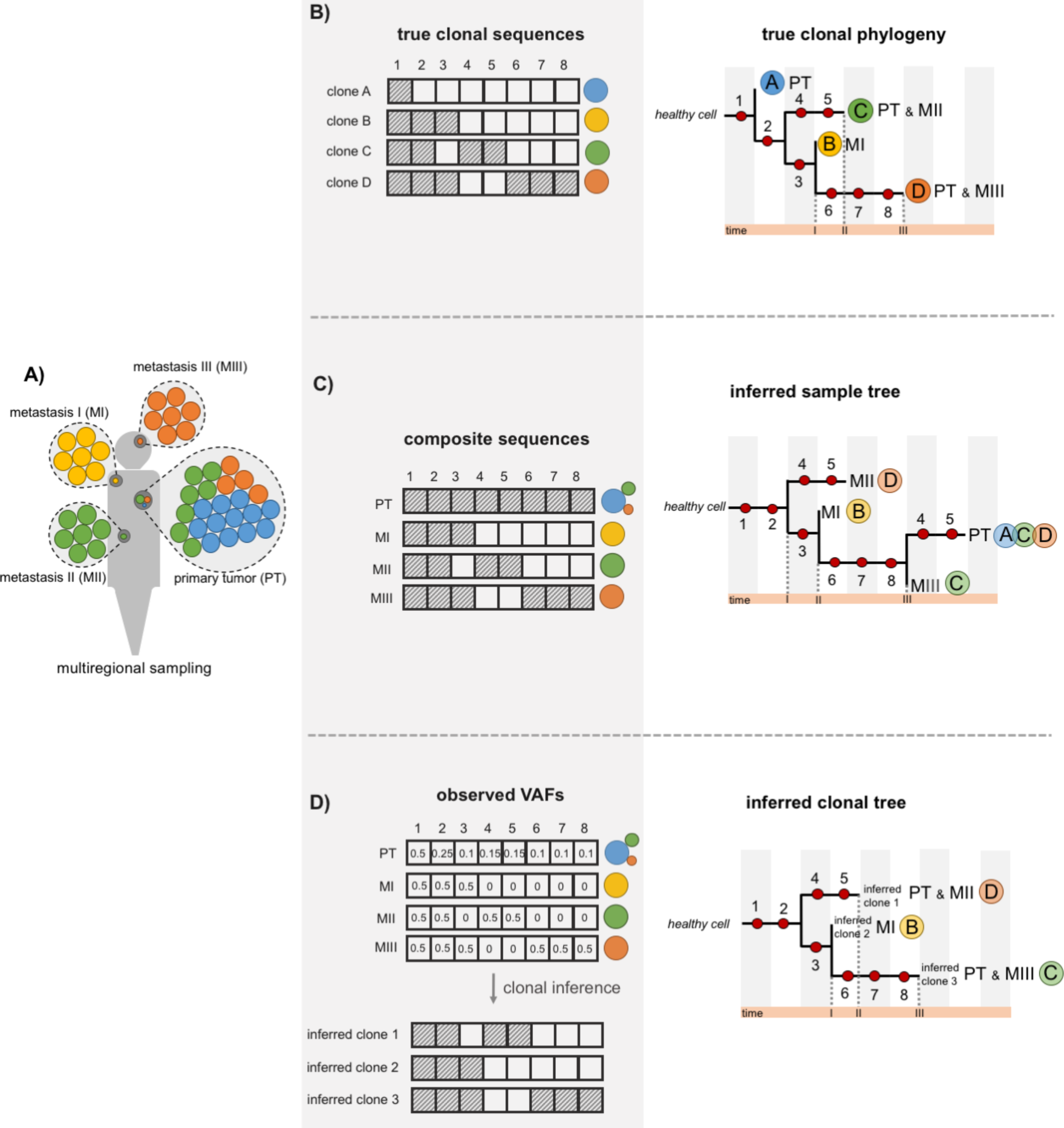
Incorrect chronological ordering of metastatic events using composite sequences. **A)** Sampling scheme of geographically distinct tumor samples: one primary tumor and three metastatic sites. Colored circles represent the four cellular clones present (i.e., A, B, C and D). **B)** *Left-panel*: Clonal sequences based on genotype information from 8 somatic mutations - dashed square indicates presence of mutation. *Right-panel*: True clonal phylogenetic tree and geographical location of each clone. Chronological order of metastatic events depicted in the orange bar below the tree. **C)** *Left-panel*: Derived regional genotype sequences using presence/absence states. *Right-panel*: Inferred sample tree using maximum likelihood or maximum parsimony for the composite sequences. Inferred chronological order of metastatic events depicted in the orange bar below the regional tree. **D)** *Left-panel*: Allele frequency estimates of each mutation per regional sample, and inferred clonal sequences using Clomial algorithm (but see supplementary note). *Right-panel*: Phylogenetic tree drawn from the inferred clones (ICs) and inferred geographical location of each clone. Inferred chronological order of metastatic events depicted in the orange bar below the clonal tree.

By assuming that a single clone occurs (or dominates) in each sample, as in Zhao*et al.*^19^, the primary tumor would be represented by a composite clone that never existed (Fig. 3C, left-panel). In consequence, if we reconstruct a MP tree from these data we will wrongly infer that metastasis II occurred before metastasis I – because in this case the lineage leading to metastasis II diverges before the lineage leading to metastasis I – and that mutations 4 and 5 evolved in parallel in the primary tumor and in metastasis II (Fig. 3C, right-panel). Moreover, because the composite clone for the primary tumor carries all mutations it is tempting to conclude that it represents the youngest clone (perhaps resulting from a recent selective sweep) when in fact is the oldest lineage. Conversely, if we use the observed VAFs to deconvolute the clones present in each sample, despite clone A not being identified by the clustering algorithm (Fig. 3D, left-panel), we will infer a phylogenetic tree that accurately represents the evolutionary history of this cancer (Fig. 3D, right-panel).

### Analysis of real data

Since the examples above represent speculative scenarios, we reanalysed three multiregional datasets in order to understand whether the use of mutational profiles *versus* the use of inferred clones can also lead to different conclusions in real scenarios. In the first study, Hao*et al.*^16^ investigated the spatial distribution of ITH in esophageal squamous cell carcinoma. Using mutational profiles, the authors reconstructed sample trees for several patients and found multiple cases where mutations were “incompatible” with the inferred tree. Interestingly, these are precisely those (parallel) mutations that appear more than once (**Supplementary Fig.2A**). Conversely, when we inferred the clones present in the samples and reconstructed their history, all parallel changes disappeared (**Supplementary Fig.2B**). We argue that in fact the latter scenario seems much more plausible.

In another study, Ling*et al.*^18^ relied on mutational profiles of 23 regional samples to investigate the evolutionary dynamics of a hepatocellular carcinoma. Unlike the original study, in which the spatial diversity patterns suggested seven major mutational lineages with well-defined spatial boundaries (**Supplementary Fig.3A**), a clonal analysis points instead to the presence of four clonal lineages segregating at different frequencies across the regional samples with substantial spatial overlap (**Supplementary Fig.3B**).

Finally, Gerlinger*et al.*^14^ explored the clonal architecture of clear cell renal carcinoma by analysing the patterns of spatial ITH in multiple patients. Although VAFs were in this case used to predict the clonal composition of each regional sample, the identification of regional subclones was limited to those samples showing strong visual evidence of multiple clonal populations (i.e., displaying clusters of mutations with clearly distinct allele frequency patterns). As a consequence, the full clonal architecture was not completely resolved, which in turn may have compromised the derived evolutionary inferences. In case EV007, for instance, clear signals of intraregional heterogeneity were only observed for two samples (R3 and R9) and the inferred MP tree suggested five instances of parallel evolution at the FAM110B, TSKU, TPRG1, NOP2 and BAP1 genes, the latter being a tumor suppressor gene identified as a putative driver (**Supplementary Fig.4A**). In contrast, a joint formal analysis of the VAFs of all regional samples suggests an alternative evolutionary scenario, with three clonal lineages showing an uneven distribution across the different subsections of the tumor (**Supplementary Fig.4B**). Notably, the clonal tree implies a single parallel mutation at the FAM110B gene.

## DISCUSSION

We have shown that the use of absence/presence mutational profiles obtained from bulk sequencing of tissue samples can compromise the study of tumor evolution. Given the pervasiveness of ITH, the types of biases we have demonstrated here - including wrong clonal histories, spurious parallel changes, reversed timings of metastases and/or incorrect phylogeographic patterns - might be commonplace suggesting that the interpretation of previous studies might need to be reevaluated. Furthermore, as already demonstrated by Kostadinov*et al.*^21^, the use of mutational profiles may also result in inaccurate branch length estimates, leading to an overestimation of substitution rate heterogeneity among samples that can be confounded with positive selection. Consequently, bulk mutational profiles need to be interpreted with caution, as without complete information of the clonal composition of each tumor different evolutionary scenarios might fit the observed ITH patterns.

Fortunately, powerful statistical inferential methods are currently being developed to characterize the clonal composition of bulk tumor samples, which generally rely on sequencing-depth information and estimates of allele frequencies^20,22-24^. While the relative performance of these methods has not been yet thoroughly benchmarked, it seems clear from our examples that they could be very helpful in reducing the level of uncertainty of the evolutionary inference from tumor bulk samples. In fact, not all multiregional tumor sequencing studies to date have relied on bulk mutational profiles. A few have already based their evolutionary inferences on clonal sequences estimated from the data (e.g., Gundem*et al.*^7^; Yates*et al.*^15^; Ding*et al.*^25^). Alternatively, single-cell sequencing data might soon become the preferred type of data for the evolutionary analysis of tumors, provided the technical limitations and the inherent sample bias arising from a limited number of cells are solved^26-28^.

Nevertheless, it is clear from our analyses that in multiregional tumor studies it is important to distinguish “sample trees”, which depict the resemblance among different regions of the tumor (or among temporal samples), from “clone trees”, which depict the history of the genetic lineages inhabiting the tumor. In evolutionary biology, these type of trees are analogous to “population/species trees” and “gene trees”, respectively (e.g., Tajima^29^, Pamilo & Nei^30^; Page & Charleston^31^ for a review). Gene trees are embedded inside population trees in the same way as clonal lineages evolve along different tumor regions, although tumor samples should be much more admixed than organismal population samples.

Outside cancer genomics, potential biases resulting from the evolutionary analysis of pooled individuals (“pool-sequencing”) have been already identified and several corrections have been proposed for the estimation of allele frequencies, diversity indices, SNP calling, population structure, tests of neutrality or association tests from pool-seq data^32^. Some of these corrections might be applicable in the cancer scenario.

In the future, complementary strategies combining clonal estimates derived from bulk-sequencing data with single-cell information should provide a more precise view of the clonal architecture of tumors, which could ultimately be used to improve cancer prognosis and therapy.

## ACKNOWLEDGMENTS

We would like to thank Andrés Pérez-Figueroa, Inigo Martincorena, Harald Detering and Sara Rocha for their comments on earlier versions of the manuscript. This work was supported by the European Research Council (ERC-617457-PHYLOCANCER awarded to D.P.) and by the Ministry of Economy and Competitiveness - MINECO (BFU2015-63774-P awarded to D.P.). T.P. was supported by a PhD fellowship from the Galician Government (ED481A-2015/083) and a PhD fellowship from the Spanish Government (FPU15/03709).

## AUTHOR CONTRIBUTIONS

D.P. conceived the project and designed the analyses. D.P., J.M.A. and T.P. performed the analyses. J.M.A. and D.P. wrote the manuscript.

